# The *Nicrophorus vespilloides* genome and methylome, a beetle with complex social behavior

**DOI:** 10.1101/023093

**Authors:** Christopher B. Cunningham, Lexiang Ji, R. Axel W. Wiberg, Jennifer Shelton, Elizabeth C. McKinney, Darren J. Parker, Richard B. Meagher, Kyle M. Benowitz, Eileen M. Roy-Zokan, Michael G. Ritchie, Susan J. Brown, Robert J. Schmitz, Allen J. Moore

**Author notes:** Corresponding Authors: CB Cunningham and AJ Moore.

## Abstract

Testing for conserved and novel mechanisms underlying phenotypic evolution requires a diversity of genomes available for comparison spanning multiple independent lineages. For example, complex social behavior in insects has been investigated primarily with eusocial lineages, nearly all of which are Hymenoptera. If conserved genomic influences on sociality do exist, we need data from a wider range of taxa that also vary in their levels of sociality. Here we present information on the genome of the subsocial beetle *Nicrophorus vespilloides*, a species long used to investigate evolutionary questions of complex social behavior. We used this genome to address two questions. First, does life history predict overlap in gene models more strongly than phylogenetic groupings? Second, like other insects with highly developed social behavior but unlike other beetles, does *N. vespilloides* have DNA methylation? We found the overlap in gene models was similar between *N. vespilloides* and all other insect groups regardless of life history. Unlike previous studies of beetles, we found strong evidence of DNA methylation, which allows this species to be used to address questions about the potential role of methylation in social behavior. The addition of this genome adds a coleopteran resource to answer questions about the evolution and mechanistic basis of sociality.

## Introduction

Understanding phenotypic evolution necessitates investigating both the ultimate and proximate influences on traits; however, these investigations require the appropriate tools. Social behavior is a particularly thorny phenotype to study because of its complexity, variation, and its multi-level integration across an organism (Boake *et al.*, 2002). In addition, social behavior also often displays unusual evolutionary dynamics arising from the genetic influences on interactions required for sociality (McGlothlin *et al.*, 2010). Although single genes can influence behavior (Fischman *et al.*, 2011), social behavior is often multifaceted and can reflect a complex genetic architecture (Walling *et al.*, 2008; Mikheyev and Linksvayer, 2015) including influences from epigenetic mechanisms (Cardoso *et al.*, 2015). Genomes in particular are useful resources for evolutionary questions of social behavior because they grant access to both broad scale and fine scale details and mechanisms (Richards, 2015). For social behavior, while there are multiple Hymenopteran genomes available to investigate fine scale detail, we lack sufficiently distantly related species to address broader patterns. It is therefore important to develop genomic resources for organisms that are particularly phenotypic models of social behavior but where genomic information is lacking.

The genomes of several social insect are now available, including eusocial species such as honey bees (Elsik *et al.*, 2014), stingless bees (Kapheim *et al.*, 2015), several ant species (Gadau *et al.*, 2012), a termite (Terrapon *et al.*, 2014), primitively eusocial species including bumble bees (Sadd *et al.*, 2015), and a sweat bee (Kocher *et al.*, 2013). However, only one, the termite (Blattodea), is outside of the insect order Hymenoptera. Although enormous progress has been made in identifying genes associated with the behavioral division of labor and developmental shifts in social and other behavioral tasks (Mikheyev and Linksvayer, 2015; Rehan and Toth, 2015), and the influence of epigenetic inheritance on developmental plasticity and behavior (Glastad *et al.* 2015; Yan *et al.*, 2015), the generality of any mechanism underlying social interactions requires information from insects reflecting other levels of sociality and from other orders. Sociality at some level occurs in nearly all insect orders (Wilson, 1971; Costa, 2006), with eusociality representing an extreme on a social continuum. Outside Hymenoptera there are many subsocial species that have highly developed social behaviors, including parental care, but no division of labor (Costa, 2006). To begin to address this gap, we assembled the genome of *Nicrophorus vespilloides*, a subsocial beetle that serves as a behavioral model species for many types of complex social interactions, including elaborate and advanced parental care with direct regurgitation of food to begging offspring (Walling *et al.* 2007), parent-offspring conflict (Kilner and Hinde, 2012), sibling competition (Smiseth *et al.*, 2007), and adult competition for resources (Hopwood *et al.*, 2013). By sequencing the genome of *N. vespilloides*, we were able to address two questions: first, is the gene complement of *N. vespilloides* more reflective of phylogeny or life history? Second, given methylation has been implicated in the success of eusocial species and facilitate plasticity, could this mechanism play a role in *N. vespilloides* social plasticity as well? *Tribolium castaneum*, the model beetle species, seems to lack DNA methylation (Zemach *et al.*, 2010). This has lead to the assumption that methylation may be unimportant in beetles generally. However, methylation has been suggested to regulate behavioral states in social insects (Glastad *et al.* 2015; Yan *et al.* 2015). *Nicrophorus vespilloides* is an unusual beetle in that it is highly social, with extensive interactions between parents and offspring, but males in the presence of females do not care for offspring or show the same levels of gene expression as caring parents (Parker *et al.*, in review). There is a rapid transition between behavioral states if the female parent is removed (Smiseth *et al.* 2005), with extensive changes in gene expression in the male (Parker *et al.*, in review). Given this, we hypothesized the presence of DNA methylation in *N. vespilloides*, which could provide a mechanism for this rapid behavioral transition.

Burying beetles (*Nicrophorus spp.*) are a group of about 70 species that have a long history as a model species used to investigate the evolution of complex social behaviors (Eggert and Müller, 1997; Scott, 1998; Sikes and Venables, 2013). Burying beetles use vertebrate carcasses as food for their offspring, but go well beyond simple forms of parental care such as providing food and protection, and display extensive and complex family interactions (Eggert and Müller, 1997; Scott 1998; Trumbo 2012). Once suitable carrion is located, these beetles remove hair, feathers, or scales, form the carcass into a ball, and spread anti-microbial excretions over its surface to retard decomposition by microbial growth. During the period of carcass preparation, the female deposits eggs in the soil around the carcass and upon hatching larvae crawl to the carcass and enter a small cavity opened by the parents in the carcass. In the next phase, there are extensive and direct parent-offspring interactions involving begging by offspring and direct regurgitation of digested carrion by parents to begging offspring. Parents also pre-digest carrion and continue to process the carcass by mastication and excreting digestive enzymes into the larval cavity to assist larvae with easy food acquisition. Parents also continue to maintain the carcass against microbial growth and interspecific competitors (e.g., fly larvae). Both parents can be present, and there is communication between parents as well as between parents and offspring. The most extensively studied burying beetle, *N. vespilloides*, has proven an excellent model for investigating the ecology and evolution of social interactions between family members (Eggert and Müller, 1997; Scott, 1998; Trumbo, 2012; fig. 1). Although parental care is essential, especially in the first 24 hours of larval life (Eggert *et al.*, 1998; Smiseth *et al.*, 2003), care in this species can be uniparental, either male or female, or biparental. The parental care expressed is identical in all three conditions, and all forms of care are equivalently beneficial for offspring (Eggert, 1990; Parker *et al.*, *in review*). Although parents attend the carcass for 95 h after larval arrival (Parker *et al.*, *in review*), there is variation among parents in how much time they allocate to care and its different components, and these differences are heritable in *N. vespilloides* (Walling *et al.*, 2008).

**Fig. 1.**
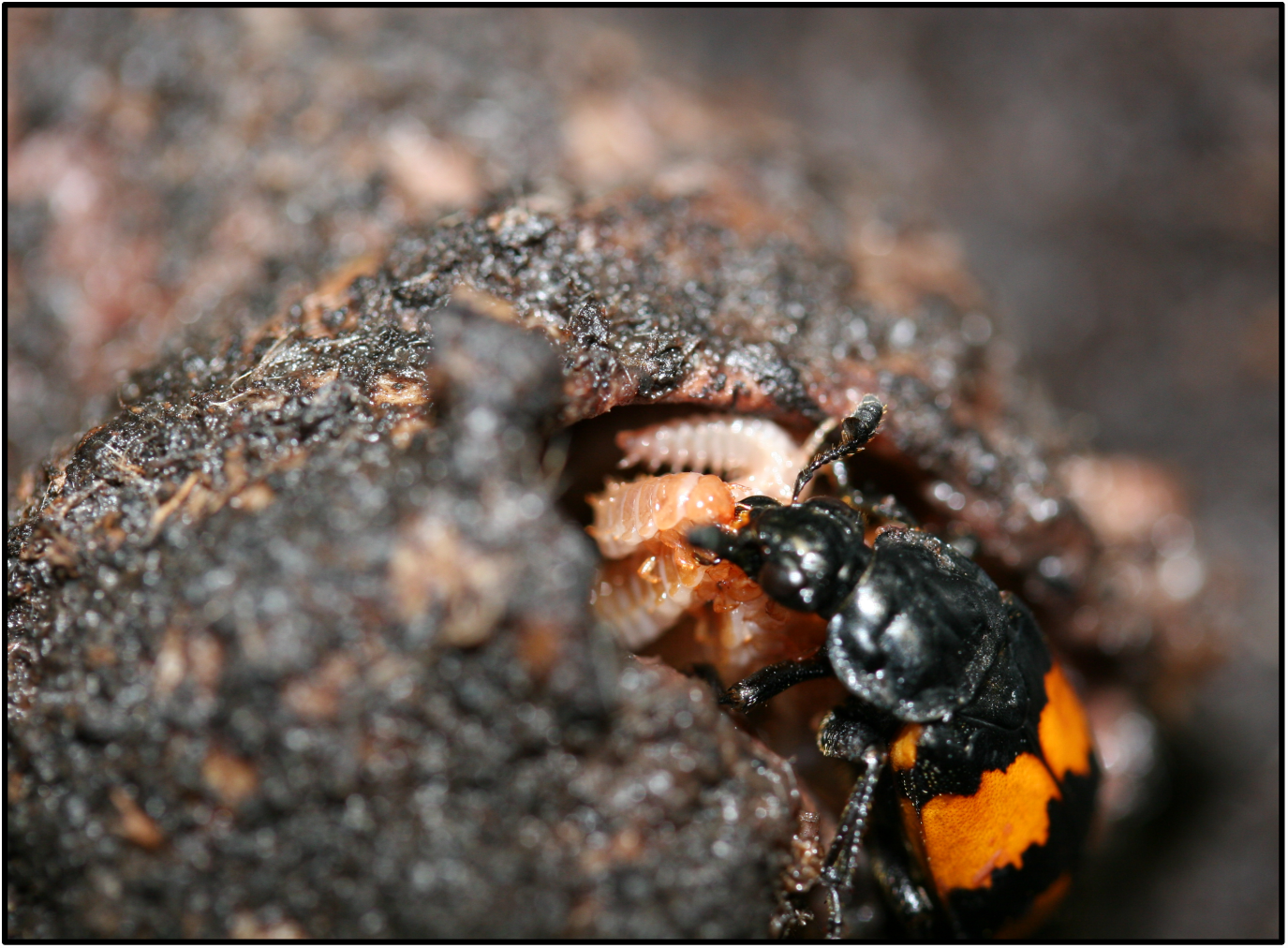
An adult female *Nicrophorus vespilloides* regurgitating food into the mouth of her begging larvae on a prepared mouse carcass. Photograph by A. J. Moore.

Here, we report a genome assembly of *N. vespilloides* and use this to investigate hypotheses regarding evolution associated with social behavior. Our assembly integrates high-throughput short reads, long reads, and a genome map providing sequence for greater than 90% of the predicted genome size. We annotated 13,526 protein-coding genes and compared these genes to social species, another beetle, and other insects that use vertebrate carcasses as food but lack sociality. The overlap of shared number of orthologs was similar between *N. vespilloides* and all other insect groups regardless of life history. We then tested if *N. vespilloides* has DNA methylation by looking for the enzymes responsible for DNA methylation and further by using whole genome bisulfite sequencing with the hypothesis that *N. vespilloides* would lack DNA methylation as does *T. castaneum* (Zemach *et al.*, 2010). Unlike previous studies of beetles, we found evidence of DNA methylation. This genome adds the first coleopteran resource to answer questions about the evolution and mechanistic basis of complex social behavior.

## Results

### Genome Sequencing and Assembly

We assembled the genome of *N. vespilloides* by integrating evidence from Illumina short reads, Pacific Bioscience (PacBio) continuous long reads, and a BioNano Genomics genome map (supplementary table S1). We assembled 195.3 Mb of the *N. vespilloides* genome, which is 95.7% of its predicted size (supplementary fig. S1). The draft genome is contained within 5,858 contigs with an N50 of 102.1 kb and further into 4,664 scaffolds with an N50 of 122.4 kb (Longest scaffold: 1.80 Mb; table 1). The GC content is 32%, consistent with two other beetle genomes, *T. castaneum* at 33% (Tribolium Genome Sequencing Consortium, 2008) and *Dendroctonus ponderosae* at 36% (Keeling *et al.*, 2013).

**Table 1.**
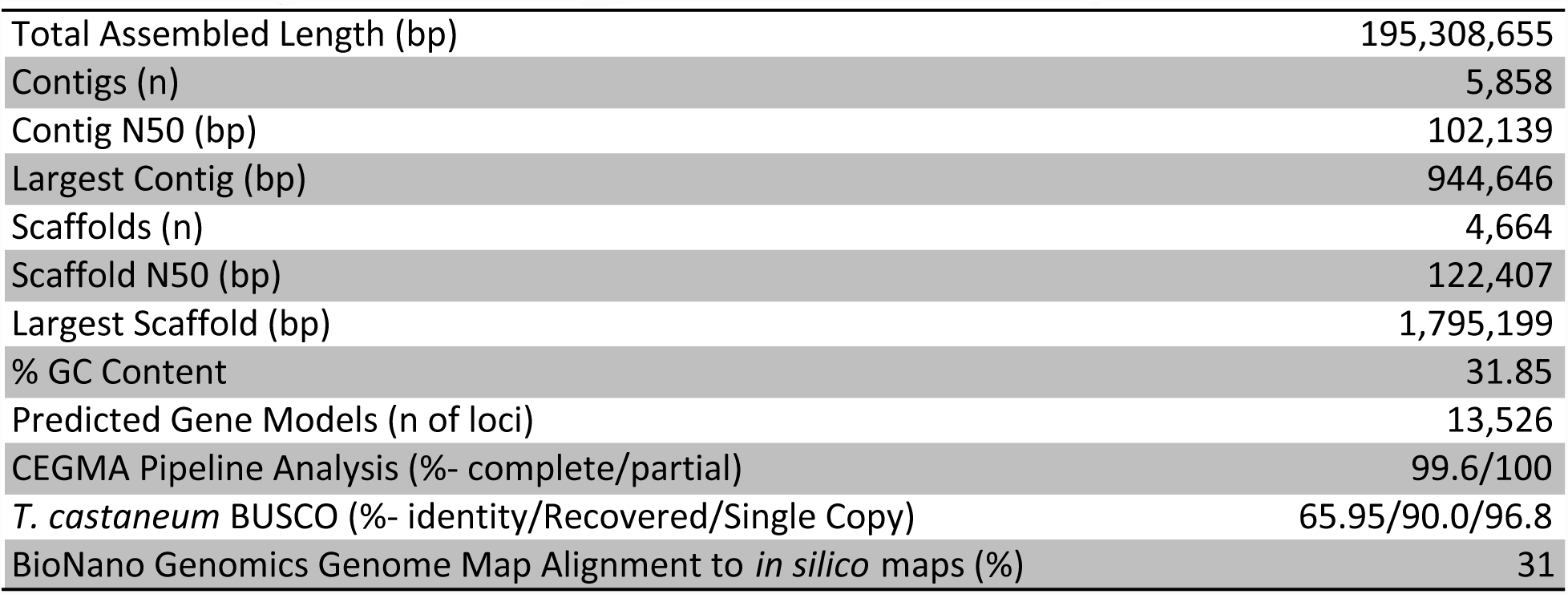
Summary Statistics of N. vespilloides Draft Genome Assembly.

We assessed how well the protein-coding portion of the genome was assembled using the CEGMA and BUSCO pipelines. Our genome contained 247 complete orthologs (99.6%) and 248 partial orthologs (100%) of the CEGMA proteins. Of the 2,827 *T. castaneum* BUSCO proteins, our genome contained 2,737 (96.8%) as single-copy orthologs and 86 (3.1%) as multi-copy orthologs. We also mapped the RNA-Seq data used for annotation back to the genome to assess transcriptome coverage. There was an 89.7% mapping rate.

### Genome Annotation

We used Maker2 to annotate the protein-coding portion of the genome by integrating *ab initio*, protein homology, and species-specific RNA-Seq evidence into consensus gene models. We obtained 13,526 predicted gene models. The gene models had an average protein length of 466.7 amino acids and 6.3 exons. Maker2 also predicted 5’ untranslated regions (UTR’s) for 5,813 genes (mean: 512 bp) and 3’ UTR’s for 4,549 genes (mean: 980 bp).

We were able to functionally annotate 11,585 gene models (85.6%) against UniProtKB with BLASTp. Restricted to species that had five or more best matches against *N. vespilloides* (encompassing 97.8% of the annotated gene models), the annotated gene models overwhelmingly returned the strongest similarity to other Coleoptera (fig. 2, supplementary table S2; top three species- *T. castaneum*: 6,969, *D. ponderosae*: 1,368, *Anoplophora glabripennis*: 743; Coleopteran total: 9,210 (79.5%)). Arthropods were the strongest similarity matches for 11,305 (99.7%) gene models (fig. 2). We were also able to identify at least one protein domain in 86.1% of the genes using InterProScan5. Searches against the Pfam database found 9,467 domains from 3,932 unique families. We were also able to assign at least one Gene Ontology (GO) term to 7,492 genes (55.4%). Additionally, we were able to associate KEGG orthology terms with 44.8% of the genes.

**Fig. 2.**
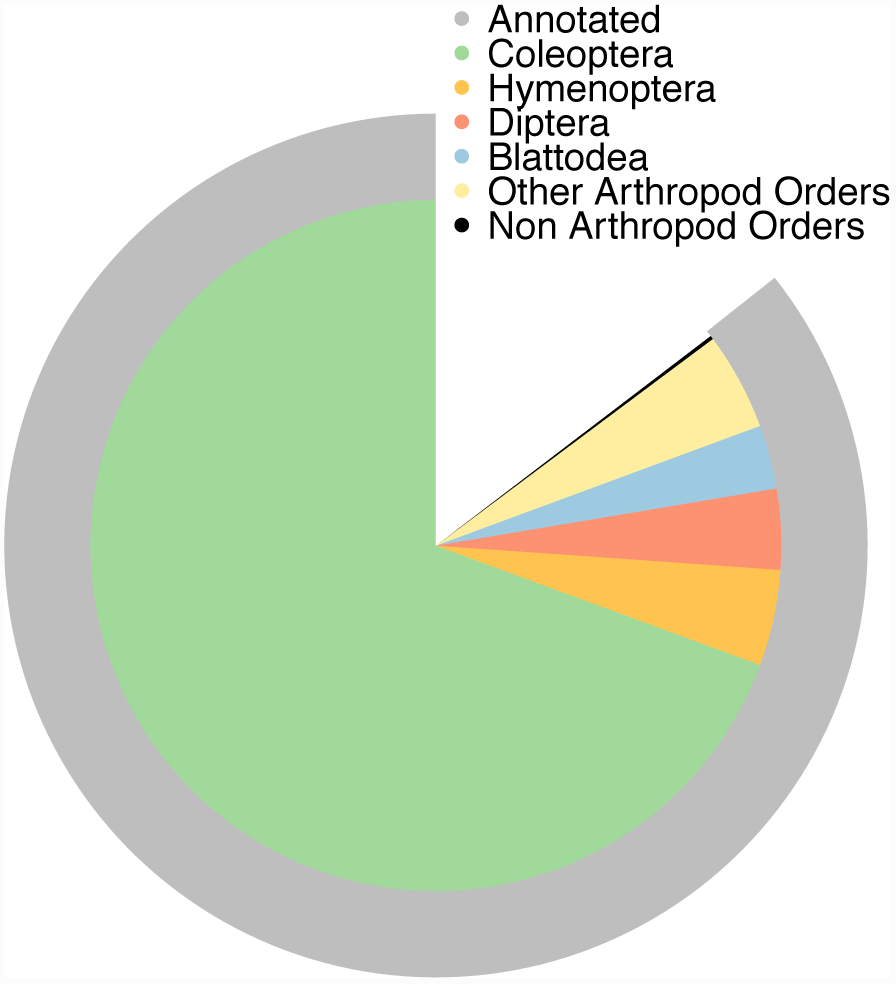
A two-ring pie chart showing results of annotation with BLAST against the complete UniProtKB database. First outer ring (grey) shows the proportion of gene models that could be annotated. Second ring (multi-colored) shows the proportion of best BLAST hits of the annotations by Order for all species with 5 or more best hits (97.8%). The best BLAST hits were overwhelmingly from other beetles and other Arthropods.

Our *de novo* repeat analysis found that 12.85% of the draft genome is composed of repetitive elements. The top three classifications of repeats were: unclassified repetitive elements (6.13%), DNA elements (3.35%), and simple repeats (2.24%). The overall repeat content is lower than that reported for beetles *T. castaneum* (Tribolium Genome Sequencing Consortium, 2008) and *D. ponderosae* (Keeling *et al.*, 2013), but higher than the honey bee (Elsik *et al.*, 2014) and the red harvester ant (Smith *et al.*, 2012), all of which have genomes that are of comparable size to *N. vespilloides*. Additionally, when we provided our repeat library to RepeatMasker to mask the *T. castaneum* genome only 1.65% was masked, an outcome consistent when the repeat library of *D. ponderosae* was used for the same task (0.15% of *T. castaneum* masked; Keeling *et al.*, 2013; supplementary table S3).

### Orthology of Gene Models

We used OrthoMCL, which clusters proteins based on a reciprocal best BLAST hit strategy, to assign orthology of the *N. vespilloides* proteome against five other insect proteomes chosen either because they are genomic models (*T. castaneum, Drosophila melanogaster*) or because of their life history (*Apis mellifera, Musca domestica, Nasonia vitripennis*). Our analysis produced 11,929 orthologous groupings with representatives from at least two different lineages. There were 4,928 orthologs groupings that contained at least one protein from each species. Of these, 3,734 groups were single copy orthologs among the six insects. There were 153 groupings containing 532 proteins that had proteins from *N. vespilloides* only. The beetles, *N. vespilloides* and *T. castaneum*, were represented in 7,827 groupings and 716 groupings were exclusive to beetles (650 were single copy ortholog groupings). We then made two specific comparisons of the proteomes of *N. vespilloides*, *T. castaneum*, *A. mellifera*, *N. vitripennis*, *D. melanogaster,* and *M. domestica* (fig 3). We first compared *N. vespilloides* to a social (*A. mellifera*) and non-social (*N. vitripennis*) Hymenopteran, including *T. castaneum* as a non-social beetle (fig. 3A). There was no increased similarity in the number of protein families between *N. vespilloides* and the Hymenoptera than there was between *T. castaneum* and the Hymenoptera. We next compared the proteome of *N. vespilloides* to a Diptera that uses rotting carcasses for reproduction, *M. domestica* and again included *T. castaneum* and another dipteran, *D. melanogaster* (fig 3B). Again, there were no obvious differences in overlap between *N. vespilloides* and the Diptera that reproduces on carrion and *T. castaneum* and *D. melanogaster*. Thus, the predicted proteins of *N. vespilloides* appear to be as similar to the other insects as is another beetle, *T. castaneum* and there is no obvious specialization associated with sociality or reproductive mode.

**Fig. 3.**
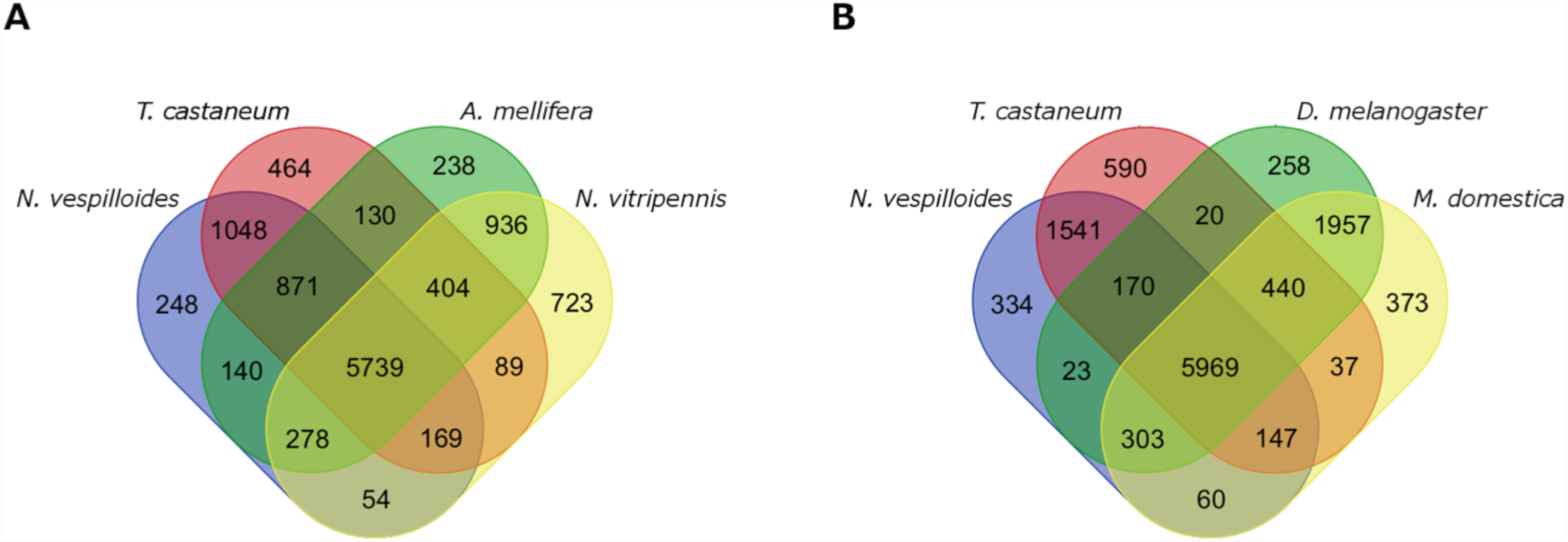
Figure shows the results of the OrthoMCL analysis that clustered the proteomes of *Nicrophorus vespilloides*, *Tribolium castanuem*, *Apis mellifera*, *Nasonia nitripennis*, *Drosophila melanogaster*, and *Musca domestica* into orthologous groupings. (A) A Venn diagram showing the overlap in the orthologous groupings of the two beetles (*T. castaneum* and *N. vespilloides*) and the two Hymenoptera (*A. mellifera* and *N. vitripennis*). (B) A Venn diagram showing the overlap in orthologous groupings of the two beetles (*T. castaneum* and *N. vespilloides*) and the two Diptera (*D. melanogaster* and *M. domestica*).

### Gene Family Expansion and Contraction

To investigate if there had been any gene family expansions or contractions in *N. vespilloides*, we analyzed the results of the OrthoMCL analysis with CAFÉ. There were 269 orthology groupings (or gene families) that showed significant expansion or contraction between the six insect species compared at *P* < 0.0001. Of these groupings 12 showed significant differences within the *N. vespilloides* lineage. There were 8 expansions and 4 contractions (supplementary file S1). The expansions were mostly families of uncharacterized proteins (7/8), while the last family was a chymotrypsin protease. There was not an enrichment of any GO term from the expanded gene families. The contracted families had highest similarity to an esterase, a transpoase, a cytochrome P450, and an uncharacterized protein in *T. castaneum*. Some of these are also differentially expressed during caring (Parker *et al.*, in review).

### Selection Analysis

Signatures of selection on the protein coding genes of *N. vespilloides* was investigated by comparing the *dN*/*dS* (*ω*) ratio to *T. castaneum* and *D. ponderosae* for the 5,584 one-to-one orthologs we detected between these lineages. Twenty five genes showed signs of differential divergent selection in the *N. vespilloides* lineage after our filtering criteria were applied (BH false discovery rate = 0.05 and removal of genes showing *dN, dS,* or *ω* >10). The distributions of these estimates are shown in fig 4. Two genes show evidence of positive selection *ω* > 1. Ephrin-B2 (*efn-b2*; *ω* = 1.45) and NK Homeobox (HOX) 7 (*nk7*; *ω* = 2.16). *efn-b2* also has a *ω* > 1 in the other lineages (*ω* = 1.5) while *nk7* shows evidence of strong conservation in the *T. castaneum* and *D. ponderosae* lineages. The median estimates of *dS, dN* and *ω* were higher in the *N. vespilloides* lineage (*N. vespilloides*- 0.0489, *T. castaneum-* 0.0487 and *D. ponderosae*- 0.0487), although not statistically significantly different.

**Fig. 4.**
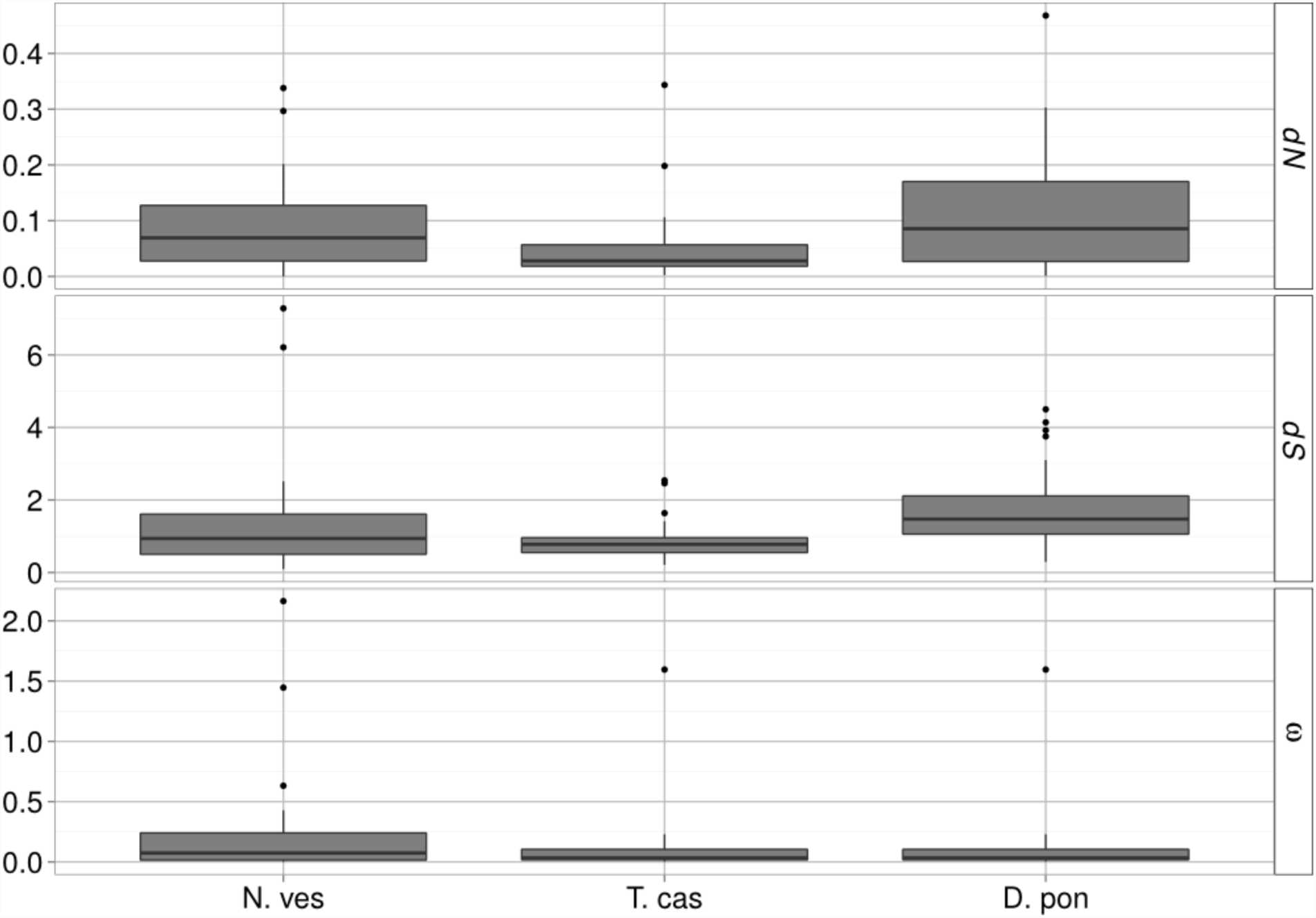
Distributions of *dN*, *dS* and *ω* for 25 genes, filtered for *dN, dS* or *ω <* 10, that show evidence of differential rates of sequence evolution on the *Nicrophorus vespilloides* (N. ves) lineage in comparison to the *Tribolium castaneum* (T. cas) and *Dendroctonus ponderosae* (D. pon) lineages.

### DNA Methylation

We used two approaches to investigate if the *N. vespilloides* genome has active DNA methylation. First, we looked for the enzymes responsible for methylation in animals (Dnmt1, Dnmt2, and Dnmt3) to determine if the machinery was present for the establishment and maintenance of DNA methylation. Second, we generated single-base resolution maps of DNA methylation using whole genome bisulfite sequencing.

Single copies of all three DNA methyltransferases were in the *N. vespilloides* genome; *T. castaneum* contains only Dnmt1 and Dnmt2 (Kim *et al.*, 2010). The methyltransferases clustered with their putative orthologs (fig. 5A). Next, using MethylC-Seq we found direct evidence for DNA cytosine methylation in *N. vespilloides* and no evidence for DNA methylation in *T. castaneum*, supporting previous reports on the latter (fig. 5B; Zemach *et al.*, 2010). Methylation (5’-methlycytosine) in *N. vespilloides* was found within a CpG context exclusively (fig. 5C). A small proportion (1.87%) of CpH (H= A,T, or C) was found during the first analysis; however, further analysis of the originally identified CpH methylated sites revealed that >98% of them were artifacts of segregating SNPs. Therefore, only strong evidence was found for CpG methylation in the genome. Methylated cytosines in *N. vespilloides* exhibited several typical insect patterns by being either reliably methylated or not (fig. 5D) and exhibited a high level of symmetrical methylation on opposing DNA strands (fig. 5E), a pattern typical of animals. The genome-wide pattern of DNA methylation observed for *N. vespilloides* is also similar to other insects. Most prominently, the majority of methylation was found in the exons (62.55 ± 0.26% of the observed methylation) and much lower levels were found in introns (10.29 ± 0.12% of the observed methylation; fig. 5F). All three biological replicates are quantitatively similar in their distribution of methylated CpG’s over gene elements (supplementary table S6). We grouped *N. vespilloides* genes as methylated or non-methylated by comparing the level of methylation of an individual gene to the average level of gene methylation found across all genes. We found 3,298 genes that were methylated significantly higher than the null expectation (fig. 5G; supplementary file S2). Following this, we performed a GO enrichment analysis on the GO terms associated with the methylated gene set. We found that translation (GO:0006412), translation factor activity/nucleic acid binding (GO:0008135), and RNA binding (GO:0003723) were significantly enriched molecular function GO terms. Cellular macromolecule metabolic process (GO:0044260), gene expression (GO:0010467), and macromolecule biosynthesis process (GO:043170) were the three most enriched biological process GO terms (see also supplementary table S7). At the level of individual genes, methylation was highest in the most exons of a gene (Fig. 5H). Methylation was also observed in the 5’- and 3’-UTR’s, with the typical steep decrease in methylation observed at the translational start site. We also observed methylation in the “promoter” region 1kb upstream from the first annotated gene element. Methylation peaks beginning at the second exon, although this is not a robust trend as methylation levels decrease to the same level of the first exon by the end of the second exon. Transposable elements were methylated to the same level as genomic intergenic background levels (3 vs. 5%, respectively).

**Fig. 5.**
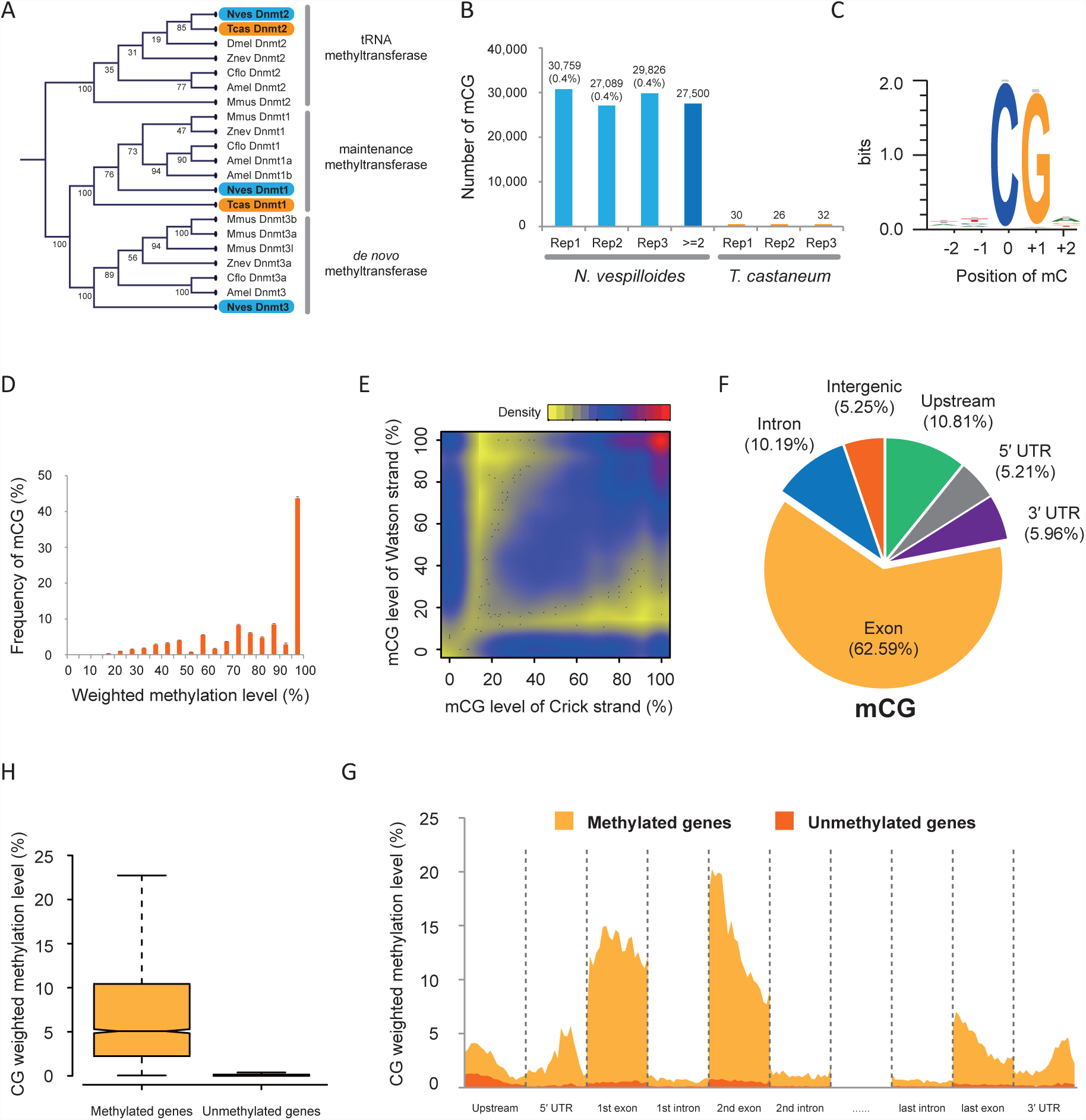
Summary of DNA methylation analyses. (A) A cladogram showing the relationship of the Dnmt’s across several insects and a mammal. Nves = *Nicrophorus vespilloides*, Tcas = *Tribolium castaneum*, Dmel = *Drosophila melanogaster*, Cflo = *Camponotus floridanus*, Amel = *Apis mellifera*, Mmus = *Mus musculus*. (B) Number of methylated cytosines in each of the three replicates of *N. vespilloides* and *T. castaneum*. (C) A sequence logo of the overwhelming occurrence of methylation in CpG dinucleotide by showing the nucleotide proportions of the two nucleotides both upstream and downstream of the methylated cytosines. (D) A histogram of the percentage of statistically methylated CpG’s versus their weighted methylation level showing the typical pattern of a cytosine either being methylated or non-methylated. (E) A density plot showing the very high symmetry of methylated CpG sites on opposing strands of DNA. (F) A pie chart showing the distribution of methylated CpG’s across gene elements. (H) A standard box plot of the methylation level of genes grouped by whether a gene was significantly more methylated compared to the average gene showing the typical bimodal where genes are either methylated or non-methylated. (G) Diagram showing the weighted methylation level of each region of a gene model summarized as 20 bins within a region. Genes were separated into “methylated” and “unmethylated” genes as in H and plotted separately.

## Discussion

The ability to detect conserved and novel molecular mechanisms that influence social behavior requires genomic resources from species across different lineages that vary in their level of sociality. Here, we report the draft genome of *N. vespilloides*, a subsocial beetle from the Silphidae. In assessing the genetic changes associated with the evolution of social behavior in insects, the *N. vespilloides* genome provides a useful line of independent evolution, offering data from outside the Hymenoptera, which diverged from Coleoptera ∼350 Mya (Wiegmann *et al.*, 2009) and at a level between solitary and eusocial. *Nicorphorus vespilloides* has sophisticated and complex parental care (Eggert and Müller, 1997; Scott, 1998; Trumbo, 2012). The highly developed social interactions between parents and offspring place this beetle at the level of “subsocial” on the evolutionary spectrum of social species (Wilson 1971; Costa 2006). Thus, we provide investigators interested in the genomic and molecular signatures of social interactions, parent-offspring conflict, social tolerance, mate choice, and mate cooperation with an experimentally tractable and evolutionarily divergent model to use in comparative studies. In addition, unlike Hymenoptera and the eusocial termites, where parental care is associated with developmental transitions to behavioral tasks, in *N. vespilloides* parental care is reversible and repeatable as individuals switch between solitary to social to solitary again across breeding cycles. Moreover, in *N. vespilloides*, both males and females express care behavior, providing a decoupling of reproductive physiology and social interactions (Roy-Zokan *et al.,* 2015). Finally, *N. vespilloides* represents an evolutionarily divergent beetle to the other available coleopteran genomes (∼240 Mya from both *T. castaneum* and *D. ponderosae;* Hunt *et al.*, 2007).

We successfully assembled the *N. vespilloides* genome using Illumina short reads, PacBio continuous long reads, and a BioNano Genomics genome map. Our assembly quality compares favorably with other recently published insect genomes; especially considering our organism is outbred (Richards and Murali, 2015). We found that our genome is similar to other recently sequenced insect genomes, with a comparable number of genes and percentage of genes having a functional annotation (Kim *et al.*, 2010; Wurm *et al.*, 2012; Keeling *et al*., 2013; Oxley *et al.*, 2014; Wang *et al.*, 2014). Our orthology analysis showed that *N. vespilloides* was as, but no more, similar to a social Hymenoptera or to Diptera that exist in rotting flesh than the asocial beetle *T. castaneum* was to either pair with respect to the number of shared gene families.

Very few of the *N. vespilloides* genes we examined showed evidence of differential rates of sequence evolution compared to the *T. castaneum* and *D. ponderosae* lineages. Among the genes that did show differential *dN/dS* ratios, the majority showed constrained evolution. We found only two genes with evidence of positive selection. NK Homeobox (HOX) 7 had an elevated *ω* in the *N. vespilloides* lineage but is highly conserved in the other lineages. Ephrin-B2 had an elevated *ω* in all lineages but it is slightly lower in the *N. vespilloides* lineage. Both of these genes are involved in patterning (Dönitz *et al.*, 2015; Dos Santos *et al.*, 2015). Overall, the genes compared show a high degree of conservation. One limitation of this analysis is the approximately 240 Mya of evolutionary distance between *N. vespilloides* and *T. castaneum*. Moving forward, it would be interesting to see how robust these results are to other types of analyses of molecular evolution and as more beetle species over a range of phylogenetic distances are available for comparison.

Beetles are often described as lacking DNA methylation, based on *T. castaneum* (Glastad *et al*, 2015; Yan *et al.* 2015). We have direct evidence for DNA methylation in the *N. vespilloides* genome. It is intriguing that a social beetle has this trait, which suggests the potential for an overlap in epigenetic mechanisms influencing social traits. Regardless, our works shows that lacking methylation is not a general feature of Coleoptera. In fact, methylation in *N. vespilloides* looks very similar to most other insects with active systems of methylation. *Nicrophorus vespilloides* has DNA methylation that is restricted to CpG sites at levels similar to honey bees (Lyko *et al.*, 2010) and the jewel wasp *Nasonia vitripennis* (Wang *et al.*, 2013), the ants *Camponotus floridanus* and *Harpegnathos saltator* (Bonasio *et al.*, 2012), a grasshopper *Schistocerca gregaria* (Falckenhayn *et al.*, 2013), a locust *Locusta migratoria* (Wang *et al.*, 2014), and the silk moth *Bombyx mori* (Xiang *et al.*, 2010). Methylation is concentrated within the exons of genes as seen with honey bees (Lyko *et al.*, 2010), ants (Bonasio *et al.*, 2012), the jewel wasp (Wang, *et al.*, 2013), but different from a locust (Wang *et al.*, 2014) and the silk moth (Bonasio *et al.*, 2012). Methylation was also found in the UTR’s, a pattern also reported in *C. floridanus* and *H. saltator* (Bonasio *et al.*, 2012). Methylation peaks at the beginning of the second exon, a pattern seen in ants (Bonasio *et al.*, 2012) and the jewel wasp (Wang *et al.*, 2013). Differential DNA methylation has been implicated in the transition between behavioral states in social insects (Lyko *et al.*, 2010; Bonasio *et al.*, 2012; Herb *et al.*, 2012; Terrapon *et al.*, 2014). Because *N. vespilloides* demonstrates dramatic and reversible switches in behavioral states across a breeding cycle, and can have multiple breeding cycles, we hypothesize that DNA methylation is an epigenetic mechanism that influences these behavioral transitions and stability.

Studies of the genetic basis and evolution of complex social behavior have focused on specific genes, with conflicting results. However, these are mostly focused on age-based division of labor in the eusocial Hymenoptera (Mikheyev and Linksvayer 2015; Rehan and Toth 2015). The addition of the *N. vespilloides* genome allows us to expand beyond hymenopteran-specific aspects of social behavior, and allows us to begin to address broader categories of social traits. Although there are numerous aspects of the life history of burying beetles that make them unique (Eggert and Müller, 1997; Scott 1998), here we have emphasized the value of using *N. vespilloides* as a model for studying family social interactions and social evolution. These beetles are particularly suited for questions of parental care because the phenotype is robust and readily measured, contains diverse sub-behaviors that are reliably observed and scored, can vary between males and females in the context in which it is expressed, and is highly replicable (Walling *et al.* 2008). With the addition of the *N. vespilloides* genome alongside Hymenoptera and a eusocial termite, we now have a taxonomically diverse arsenal of phenotypically overlapping organisms to look for phylogenetically independent genomic mechanisms and signatures of evolution, conservation, and novelty.

## Materials and Methods

### Animals Samples

All *Nicrophorus vespilloides* used in this research were obtained from an outbred colony maintained at the University of Georgia under laboratory conditions for this species (see Cunningham *et al.*, 2014 for a full description of conditions).

### Genome Size Estimation, Sequencing, Assembly, and Quality Control

We used flow cytometry with propidium iodide staining to estimate the genome size of *N. vespilloides* using *T. castaneum* as a standard. Nuclei from insect heads and whole insects, respectively, were prepared as described in Yu *et al.* (2015), stained as in Hare and Johnston (2011), and analyzed with a CyAn Flow Cytometer (Beckman Coulter, Brea, CA) at the UGA’s Center for Tropical and Emerging Global Diseases Flow Cytometry Core Facility. Data was processed with FlowJo software (Treestar, Inc., Ashland, Oregon).

Genomic DNA was extracted from a single larva derived from a single sibling-sibling mating using a SDS-lysis buffer and a phenol-chloroform extraction. A 275 bp Illumina (San Diego, CA, USA) TruSeq library was prepared and run on one lane of an Illumina HiSeq 2000 using a paired-end (2 × 100 bp) sequencing protocol at the HudsonAlpha Genome Sequencing Center (Huntsville, AL, USA).

We used FastQC (v0.11.2; Babraham Institute; default settings) to create summary statistics and to identify possible adapter contamination of raw Illumina paired end reads. No adapter contamination was reported, a result supported by analysis with CutAdapt (v1.2.1; Martin, 2011), which only found evidence for adapters in less than 0.01% of the raw reads. Because sequencing library construction can generate inserts of genomic DNA that are less than twice the average read length, overlapping paired-end reads were first merged using FLASH (v1.2.4; Magoc and Salzberg, 2011; default settings, insert size: 278 bp with SD of 53 bp (estimate from Platanus scaffolding step)). Quality control was performed with PrinSeq (Schmieder and Edwards, 2011a). Reads were required to have a mean overall Phred quality score of ≥ 25, read ends were trimmed to >20 Phred quality score, a minimum length of 90 bp and a maximum length of 99 bp were allowed, and reads were allowed only one unidentified (N) nucleotide per read.

To obtain Pacific Bioscience (PacBio; Menlo Park, CA, USA) continuous long read, we extracted genomic DNA using the same phenol-chloroform extraction as used to extract gDNA for the Illumina sequencing from a brother/sister pair of adult beetles that had been inbred for 6 generations. A 14.4kb long insert PacBio library was prepared by the University of Maryland Institute for Genomic Sciences. This library was sequenced with 22 PacBio’s RS II P5-C4 SMRT cells to generate continuous long read (CLR’s) to scaffold our assembly with to increase long-range connectivity of the genome. PacBio reads greater than 6300 bp (36.4x coverage) were error corrected with the PBcR pipeline (Koren *et al.*, 2012) using 49x coverage of the quality-controlled Illumina reads with default settings, which after error correction and assembly produced an estimated 20.9x coverage of CLRs.

To increase the long-range scaffolding (i.e., super-scaffold) of our draft genome, we generated a BioNano Genomics (San Diego, CA, USA) genome map. High molecular weight (HMW) genomic DNA was extracted from a single pupa as previously described (Shelton et al., 2015). HMW gDNA was nicked with nicking restriction digest by BspQI and BbvCI restriction enzyme that had been converted to nickases (New England Biolabs, Ipswich, MA, USA). Restriction sites were labeled with fluorescent nucleotides and imaged on the Irys system (BioNano Genomics) according to the manufacturer’s instructions.

All Illumina read passing quality control were used as input for the Platanus assembler (v1.2.1; Kajitani *et al.*, 2014). First, reads were assembled into contigs using Platanus’s assemble protocol (non-default settings: -s 3 -u 0.2 -d 0.3 -m 128). Next, contigs were scaffolded using Platanus’s scaffold protocol (non-default settings: -u 0.2). This step was iterated a total of five times using the same settings to extend the scaffold as much as possible with the Illumina reads. Gaps in the assembly were filled using Platanus’s gap_close protocol with default settings. This step was iterated twice. Only contigs/scaffolds 1kb or greater in length were used for further analysis and assembly.

PacBio reads were used to gap fill and scaffold the Platanus assembly with PBJelly2 (v14.9.9; English *et al.*, 2012) using default settings and the error corrected PacBio CLR’s reads.

A genome map created from BioNano Genomic single molecule maps was used to super-scaffold the Platanus/PBJelly assembly (Shelton *et al.*, 2015). Briefly, the images were assembled into a consensus map based on the labeling pattern of each molecule imaged. These *in silico* maps, with a cumulative length of 133.7 Mb, were compared to the predicted labeling pattern of the Platanus/PBJelly that passed a quality filter (length > 150kb and number of labels >= 8) to further scaffold and orient the Platanus/PBJelly assembly.

DeconSeq (v0.4.2; Schmieder and Edwards, 2011b) was used to assess our draft assembly for possible contamination. Besides the 1,126 bacterial species included in the distribution, we also updated the human genome sequence (h37) and added the genomes of *Caenorhabditis elegans*, *Ralstonia pickettii*, *Yarrowia lipolytica*. *C. elegans* was included because it is the closest genome available to the nematode symbiont of *N. vespilloides*, *Rhabditis stammeri* (Richter, 1993). *R. pickettii* and *Y. lipolytica* were included because they were two species that showed up when the RNA-Seq experiment was assessed for possible contamination (Parker *et al.*, in review). *Tribolium castaneum* was used as a retention database. Only one contig was flagged and removed during our contamination search; belonging to *Morganella morganii*, a common bacterium found in vertebrate intestinal tracts.

Genome assembly quality and completeness were assessed with multiple benchmark datasets. First, the CEGMA analysis pipeline (v2.4.010312; Parra *et al.*, 2009) was used to assess the completeness of 248 ultra conserved eukaryotic genes within our assembly. Next, we used the *T. castaneum* set of Benchmarking sets of Universal Single-Copy Orthologs (BUSCO’s; 2,787 genes) to further assess the assembly completeness (Waterhouse *et al.*, 2013). We also mapped the RNA-Seq reads back to the assembly to estimate coverage of the transcriptome of our assembly using the TopHat (v2.0.13) pipeline with Bowtie2 (v2.2.3) as the read aligner.

### Genome Annotation

To begin genome annotation, we first generated a *de novo* library of repeats using Repeat-Modeler (v1.0.8) that integrates three separate repeat finder programs; RECON (v1.08; Bao and Eddy, 2002), RepeatScout (v1.05; Price *et al.*, 2005), and TRF (v4.07b; Benson, 1999) with default parameters. Because some gene fragments, especially low-complexity motifs, might be captured in the repeat analysis, we used BLASTx to remove any matches to *T. castaneum* proteins in the UniProtKB database (Wang *et al.*, 2008; Jiang, 2014). The repeat analysis of the *T. castaneum*’s genome was done with RepeatMasker (v4.0.5; Smit *et al.*, 2015) using default settings.

We annotated putative protein coding genes using the Maker2 annotation pipeline (v2.31.7; Holt and Yandell. 2011) using an iterative process. After masking putative repeats within a genome, this pipeline generates gene models, including 5’ and 3’ UTR’s, by integrating *ab initio* gene predictions with aligned transcript and protein evidence. First, we mapped and assembled transcripts using the RNA-Seq data from an experiment of *N. vespilloides* in multiple behavioral states over a breeding cycle (mated, caring, and post-caring; see Parker *et al.*, 2015 for full details) using the Bowtie (v2.2.3)/TopHat (v2.0.13)/Cufflinks (v2.2.1) pipeline (Langmead *et al.*, 2010; Trapnell *et al.*, 2010; Kim *et al.*, 2013). To begin the annotation process, we annotated the genome exclusively with the *N. vespilloides* Cufflinks-assembled transcripts and the proteomes from five insects (*T. castaneum, N. vitripennis, A. mellifera, M. domestica, D. melanogaster*; downloaded from UniProtKB, including all isoforms for comprehensive coverage) using default parameters, except for est2genome=1, protein2genome=1. After this first iteration of annotation (and every subsequent iteration), three scaffolds were inspected to visually check for annotation biases (Hoff and Stanke, 2015) using Apollo genome browser (Lewis *et al.*, 2002). The next iteration used the same input data and parameters, except changes to split_hit=2000, correct_est_fusion=1, which corrected for the smaller intron size observed and the propensity of MAKER to fuse gene models that likely should be separate as inferred by visual inspection of BLAST evidence. For the next iteration, three *ab initio* gene predictors were included in the annotation process; Augustus (v2.5.5; Stanke et a., 2006), GeneMark-ES (v4.21; Lomsadze *et al.*, 2005), SNAP (v2010-7-28; Korf, 2004; using est2genome=0, protein2genome=0). With AUGUSTUS, we used the “tribolium” gene set provided with its distribution to guide gene predictions. GeneMark was trained on the draft assembly of the *N. vespilloides* genome sequence using its automated training routine. SNAP was trained using the MAKER2 gene models produced during the first round of annotation. All gene predictors were run with default parameter values. The annotation was iterated twice with the gene predictors, updating the SNAP HMM’s between the two iterations. tRNA’s were identified using tRNAscan-SE (v1.23; Lowe and Eddy, 1997) within the Maker2 pipeline during the last iteration. Other non-coding RNA (rRNA, miRNA, snRNA, snoRNA) were predicted and annotated with INFERNAL (v1.1.1; Nawrocki and Eddy, 2013) using the complete Rfam database (v12.0; Daub *et al.*, 2014; Supplementary Table S4).

### Functional Annotation of Predicted Protein-Coding Genes

To gain insight into the putative function of each gene model, we annotated our gene models with three pipelines. First, we used BLASTp (v2.2.26; Altschul *et al.*, 1997) to find the best hit against the entire UniProtKB database (vJan15; E-value: 10e-5). Next, we used InterProScan (v5.8-49.0; Hunter *et al.*, 2009) to find the known protein domains within every gene model from the TIGRFAM, ProDom, SMART, TMHMM, Phobius, PANTHER, PrositeProfiles, SignalP-EUK, SuperFamily, PRINTS, Gene3d, PIRSF, Pfam, and Coils databases. We also used InterProScan5 to assign Gene Ontology (GO) terms to further characterize each protein. KEGG pathway analysis was also performed using the KEGG Automatic Annotation Sever (KAAS; Moriya *et al.*, 2007) using the single-directional best hit method to assign orthology with default parameters and the default Eukaryote gene sets plus all available arthropod gene sets.

### Ortholog Comparison

To compare the orthology of our gene models to other insects, we analyzed our final MAKER2 proteome using OrthoMCL (v2.0.9; Li *et al.*, 2003) against five other insect proteomes (*Tribolium castaneum, Nasonia vitripennis, Apis mellifera, Musca domestica, Drosophila melanogaster*). If a gene was represented by more than one isoform in its respective OGS, the longest isoform was chosen for this analysis. We used BLASTp (E-value: 1e-5) to characterize the homology amongst all proteins. The output from this analysis was used by OrthoMCL to cluster proteins into orthologous groupings. Results are presented as Venn diagrams generated using the University of Ghent Bioinformatics Evolutionary Genomics’ Venn Diagram webtool (http://bioinformatics.psb.ugent.be/webtools/Venn/).

### Gene Family Expansion/Contraction Analysis

To investigate possible expansion and contraction of shared gene families of the six insects that we used in the OrthoMCL analysis, we used CAFÉ (v3.1; default settings; Han *et al.*, 2013) with phylogenetic relationships from Trautwein *et al.* (2012) and divergence times from TimeTree (Hedges *et al.*, 2006). Only gene families with at least one representative from *N. vespilloides* were considered as gene family contractions.

Enrichment of gene ontology (GO) terms among the expanded gene family members was performed using argiGO’s web-based Singular Enrichment Analysis (Du *et al.*, 2010) of customized annotations by comparing the GO terms associated with methylated gene from the InterProScan results to all GO terms associated with all genes from InterProScan. Specifically, a hypergeometric test with a Benjamini-Hochberg false discovery rate correction at a family-wise error rate of 0.05 was applied after GO terms were converted into generic GO slim terms. All other parameters were set at default values.

### Selection Analysis

To assess the rates of molecular evolution within the *N. vespilloides* genome, we used PAML (Yang & Bielawski, 2000; Yang, 2007) to calculate *dN, dS,* and their ratio (*w*) and compare these metrics to the *T. castaneum* and *D. ponderosae* lineages. We identified a set of 1-1 orthologs between *N. vespilloides, D. ponderosae* and *T. castaneum* using a combination of the BLAST (Altschul *et al.*, 1990; Camacho *et al.*, 2009), orthAgogue (Ekseth *et al.*, 2014), and mcl (Enright *et al.*, 2002; van Dongen, 2008) as well as part of the OrthoMCL (Li *et al.*, 2003) pipeline. In total, 5,584 orthologs between all three species were recovered. Amino acid sequences for each were aligned in PRANK (v100802; Löytynoja & Goldman, 2005). Codeml in the PAML package was used to test different models of molecular evolution for each gene. Our interest is in determining which genes show evidence of a differential rate of evolution within *N. vespilloides*. We therefore tested a basic model (model = 0, NSsites = 0, fix_omega = 0) that assumes a single *w* across all the entire phylogeny against a branch model (model = 2, NSsites = 0, fix_omega = 0), which assumes one *w* for the *N. vespilloides* branch and another *w* for the branches to *T. castaneum* and *D. ponderosae*. These models are compared, for each gene, with a likelihood ratio test with one degree of freedom. We then adjusted the significance threshold for a gene to show statistically significant different rates of sequence evolution using a Benjamini-Hochberg false discovery rate (FDR) correction at *q* of 0.05 (Benjamini & Hochberg, 1995). Finally, any estimates of *dS, dN* or *w* > 10 were discarded. These species are phylogenetically distant (240 Mya) and this increases the likelihood signals of molecular evolution will be lost due to saturation of *dS*.

### DNA Methylation Analysis

As the first step to characterize if DNA methylation existed within *N. vespilloides*, we use BLASTp (Altschul *et al.*, 1997) to identify putative DNA methyltransferases. We search our genome with known members of Dnmt families of both vertebrate (*Mus musculus*; 1, 2, 3a, 3b, 3l) and invertebrate (*T. castaneum, A. mellifera, D. melanogaster*; 1, 2, and 3). After three putative loci were found (one member per Dnmt family), we further characterized the possible functional relationship of the proteins by clustering them with the BLAST query proteins and several more invertebrate species (*Zootermopsis nevadensis* and *Camponotus flordanus*) using ClustalW followed by a Neighbor Joining tree with 10,000 bootstraps in CLC Sequence Viewer (v7.5; http://www.clcbio.com) with default settings.

To address if DNA methylation is present in *N. vespilloides* at all, we performed methylC-Seq (Lister *et al.*, 2008), whole genome sequencing on bisulfite treated genomic DNA, on three biological replicates of whole larvae to create single base resolution of DNA methylation, if present. DNA was extracted from three whole *N. vespilloides* larvae respectively using the same protocol as for the Illumina and PacBio sequencing (see above). Due to previous reports that *T. castaneum* contains no DNA cytosine methylation (Zemach *et al.* 2010), samples from this species were used as negative control and DNA was extracted from three biological replicates that each contained at least 15 pooled whole larvae using the same protocol as for *N. vespilloides*. MethylC-Seq libraries were prepared according to the following the protocol of Urich *et al.* (2015). Deep sequencing was performed using an Illumina NextSeq500 Instrument at the University of Georgia Genomics Facility (Supplementary Table S5).

Raw fastq files were trimmed for adapters CutAdapt (v1.3) and preprocessed to remove low quality reads. We aligned quality controlled reads to the *N. vespilloides* v1 and *T. castaneum* v3.0 reference genomes, respectively, using the same method as previously described in (Schmitz *et al.*, 2013). The *T. castaneum* genome and OGS gff (v3) were obtained from BeetleBase.org. Lambda sequence (which is fully unmethylated) was used as control to calculate the efficiency of the sodium bisulfite reaction and the associated non-conversion rate of unmodified cytosines, which ranged from 0.10-0.11% (Supplementary Table S5). Only cytosine sites with a minimum coverage of at least three reads were used for subsequent analysis. A binomial test coupled with Benjamini-Hochberg correction was adopted to determine the methylation status of each cytosine. Weighted methylation levels were calculated as previously described (Schultz *et al.*, 2012).

*Nicrophorus vespilloides* replicate 1 was used to compute the exact values and percentages and plot Figure 5C, 5E, 5F, 5H, and 5G, but all replicates were qualitatively similar (Supplementary Table S7). Methylated cytosines and their flanking two bases were extracted out for sequence conservation analysis using the program WebLogo 3.3 (Crooks *et al.*, 2004). To perform symmetry analysis, both strands of each CpG dinucleotide were required to have a minimum coverage of at least three reads and at least one of the CpG sites was identified as methylated. Upstream regions were defined as 1kb upstream starting from the translational start site or the transcriptional if a 5’ UTR was annotated. The program bedtools was used to determine the distribution of methylated CpG sites (Quinlan and Hall, 2010). To identify “methylated” versus “unmethylated” genes, a binomial test was performed on genes that contained at least five methylated CpG sites. The weighted methylation level (total methylated cytosines / total mapped cytosines within a region of interest) was determined for all genes that had at least five CpG sites and this was used as the probability of a gene being methylated. These results were then corrected for multiple testing using Benjamini-Hochberg correction with an FDR cut-off of 5% (Benjamini and Hochberg, 1995). For figure 5G, each gene element was divided evenly into 20 bins and the weighted methylation levels were calculated for each bin.

Enrichment of gene ontology (GO) terms among the methylated genes was tested as above.

All analyses were conducted on the zcluster housed at the Georgia Advanced Computing Resource Center at the University of Georgia.

## Acknowledgements

We would like to thank the Quantitative Biology Consulting Group at the University of Georgia (especially, Walter Lorenz and Saravanaraj Ayyampalayam), Julie Brown at the University of Georgia’s CTEGD Cytometry Shared Resource, Jessica Kissinger, the Genomic Services Lab at HudsonAlpha, Roger Nilsen at the Georgia Genomics Facility, the Institute for Genome Science at the University of Maryland, and the staff at the Georgia Advanced Computing Resource Center (especially, Yecheng Huang). Special thanks to Nick Talbot, who first suggested to AJM he develop a genome from short reads in 2007.

All genomic data has been submitted to public repositories and can be found using the NCBI BioProject number PRJNA284849.

This work was supported by the University of Georgia’s Office of the Vice-President for Research to AJM and RJS and an National Science Foundation grant (IOS-1354358) to AJM. No funding agency was involved with the design or execution of this research.

